# MDA-LDL vaccination induces athero-protective germinal center-derived antibody responses

**DOI:** 10.1101/2021.10.25.465690

**Authors:** Inmaculada Martos-Folgado, Cristina Lorenzo, Christian E. Busse, Pilar Delgado, Sonia Mur, Laura Cobos, Jose Luis Martín-Ventura, Joan Carles Escolà-Gil, Hedda Wardemann, Almudena R Ramiro

**Affiliations:** B Lymphocyte Biology Lab, Centro Nacional de Investigaciones Cardiovasculares (CNIC), Madrid, Spain; Division of B Cell Immunology, German Cancer Research Center, Heidelberg, Germany; Centro de Biología Molecular Severo Ochoa, Consejo Superior de Investigaciones Científicas, Universidad Autónoma de Madrid, Madrid, Spain; Unidad de Presentación y Regulación Inmunes, Instituto de Salud Carlos III, Madrid, Spain; CIBER de Enfermedades Cardiovasculares (CIBERCV), Madrid, Spain; Vascular Pathology Lab, IIS-Fundación Jiménez Díaz-Universidad Autónoma, Madrid, Spain; CIBER de Diabetes y Enfermedades Metabólicas Asociadas (CIBERDEM), Madrid Spain; Institut de Recerca de l’Hospital de La Santa Creu I Sant Pau, Barcelona, Spain

## Abstract

Atherosclerosis is a chronic inflammatory disease of the arteries that can lead to thrombosis, infarction and stroke and is the leading cause of mortality worldwide. Immunization of pro-atherogenic mice with malondialdehyde-modified low-density lipoprotein (MDA-LDL) neo-antigen is athero-protective. However, the immune response to MDA-LDL and the mechanisms responsible for this athero-protection are not completely understood. Here, we find that immunization of mice with MDA-LDL elicits memory B cells, plasma cells, and switched anti-MDA-LDL antibodies as well as clonal expansion and affinity maturation, indicating that MDA-LDL triggers a *bona fide* germinal center antibody response. Further, *Prdm1*^fl/fl^ *Aicda*-Cre^+/ki^ *Ldlr*^-/-^ pro-atherogenic chimeras, which lack germinal center-derived plasma cells, show accelerated atherosclerosis. Finally, we show that MDA-LDL immunization is not athero-protective in mice lacking germinal-center derived plasma cells. Our findings give further support to the development of MDA-LDL-based vaccines for the prevention or treatment of atherosclerosis.

## INTRODUCTION

Cardiovascular disease (CVD) remains the leading cause of mortality in the world, with most CVD deaths resulting from myocardial infarction and stroke. The main cause underlying thrombosis and cardiovascular (CV) events is atherosclerosis, a chronic disorder of large arteries caused by hyperlipidemia that leads to atheroma plaque formation ^1–5^. Thus, there is a critical need to expand and improve our therapeutic and prognostic avenues to atherosclerosis ^5,6^.

Atherosclerosis initiates with the infiltration and retention of cholesterol-rich LDL particles in the subendothelial space of the artery wall ^1,7^. Retained LDL is subject to oxidation, which leads to diverse oxidized forms of LDL (OxLDL), including MDA-LDL ^4,8^. OxLDL particles activate endothelial cells and induces the expression of adhesion molecules and chemokines that recruit monocytes to the intima layer and promote their differentiation into macrophages. At the intima, macrophages internalize OxLDL and eventually give rise to lipid-laden foam cells, which contribute to plaque growth and the generation of a necrotic core ^5,9,10^.

Adaptive immunity contributes in various and intertwined ways to atherosclerosis modulation. Different subsets of T cells have been ascribed different roles in atherosclerosis, which generally seem to stem from their pro- or anti-inflammatory properties ^11–15^. In the case of B cells, detailed studies on the contribution of different B cell subsets to atherosclerosis have revealed a complex landscape. B1 cells, a minor subset of B cells of fetal origin that produce natural IgM antibodies, are generally believed to be athero-protective ^16–18^. Marginal zone B cells are splenic non-circulating B cells which mediate an innate-like rapid antibody response to blood-borne antigens and have athero-protective capabilities^14^. Follicular B cells are essential for long-lasting, T-dependent antibody responses and have been assigned both pro-atherogenic and athero-protective roles^19–25^.

Upon antigen encounter and in the presence of T-cell help, follicular B cells became activated and can engage into the germinal center (GC) reaction and give rise to long-lived memory B cells (MBC) and high-affinity plasma cells (PC) with alternate isotypes. Key to these antibody responses are the Class Switch Recombination (CSR) and the Somatic Hypermutation (SHM) reactions. CSR is a recombination reaction that exchanges the primary IgM constant region by a downstream constant region, generating IgG, IgA or IgE isotypes. SHM introduces changes in the antigen-recognizing, variable region of antibody genes thus giving rise to clonal variants. Coupled to affinity maturation, SHM allows the generation of higher affinity antibodies. Thus, GCs are essential for high affinity humoral responses and underlie the protective mechanism of many vaccination strategies. Disruption of T-B cell interactions, abrogation of the GC reaction or broad depletion of PCs resulted in smaller plaques ^26^, but they showed more instability ^27^. On the other hand, pro-atherogenic mice where antibody secretion was abolished had larger plaques ^28^. These results reveal a complex role of the antibody response in atherosclerosis. Of note, the role of antibody responses after specific challenge with atherosclerosis antigens was not addressed in these studies.

Indeed, very few antigens have been associated with atherosclerosis. The atherosclerosis adaptive immune response is thought to be triggered by self- and neoantigens generated during disease development, the paradigm of which are oxidation-specific epitopes present in OxLDL ^6^. Heat shock protein (Hsp) 60, a mitochondrial chaperone, has also been associated with atherosclerosis ^29–33^. In addition, the aldehyde dehydrogenase 4 family member A1 (ALDH4A1) mitochondrial protein has been recently identified as an auto-antigen in atherosclerosis ^34^. Circulating ALDH4A1 is increased in mice and humans with atherosclerosis and anti-ALDH4A1 antibody is athero-protective ^34^. However, LDL forms have been by far the most studied atherosclerosis antigen and as of today are considered the best antigenic target for the design of atherosclerosis vaccines ^6^. T cells, B cells and anti-OxLDL antibodies have been found in atheroma plaques and in plasma from atherogenic mice, rabbits and humans ^35–41^.

Malondialdehyde (MDA) forms covalent adducts with lysine residues of the apoB-100 protein and generates MDA-LDL, the predominant form of OxLDL ^42,43^. MDA-LDL immunization reduces atherosclerosis ^44–48^, but the immune response to MDA-LDL is not completely understood and several mechanisms have been proposed to explain MDA-LDL-mediated atheroprotection ^12,49–51^.

Here, we find that athero-protective MDA-LDL immunization triggers an antibody immune response that entails GC formation, and the generation of MBC and PC. A high - throughput single-cell analysis of the immunoglobulin repertoire in MDA-LDL immunized mice revealed clonal expansion and accumulation of SHM, featuring a GC response. Using *Prdm1*^fl/fl^ *Aicda*-Cre^+/ki^ *Ldlr*^-/-^ chimeras, we futher show that depletion of GC-derived PCs aggravates atherosclerosis, indicating a global athero-protective role of these cells. Finally, we found that athero-protection by MDA-LDL immunization is impaired in *Prdm1*^fl/fl^ *Aicda*-Cre^+/ki^ *Ldlr*^-/-^ chimeras. Together, our results indicate that MDA-LDL immunization triggers a bona fide GC reaction and the generation of GC-derived PCs, which contribute to athero-protection.

## RESULTS

### Athero-protective MDA-LDL immunization induces GC reactions

To gain insights into the impact of MDA-LDL immunization on atherosclerosis, we immunized *Ldlr*^-/-^ mice with MDA-LDL in complete Freund’s adjuvant (CFA) or with CFA alone as described before (Figure 1A)^52^. We did not detect significant differences in weight gain or plasma lipids between groups (Figure S1A, Table S1). Quantification of atherosclerotic plaques in aortic roots showed a significant decrease in atherosclerosis extension in mice immunized with MDA-LDL+CFA compared to those immunized with CFA alone (Figure 1B-D), as previously described ^44,45,52,53^. We did not detect changes in the proportions of GC B cells or total PCs (Figure S1B-C). However, the proportion of MBCs was significantly increased in MDA-LDL+CFA immunized mice compared to CFA controls (Figure 1E). Likewise, the proportion of IgG1+ PCs was much larger in MDA-LDL+CFA mice (Figure 1F). Indeed, IgM, IgG1, IgG2b and IgG2c MDA-LDL specific antibodies were greatly increased in serum from MDA-LDL immunized mice (Figure 1G). Thus, MDA-LDL immunization of *Ldlr*^-/-^ mice is athero-protective and promotes B cell response that generates MBCs, switched PCs and MDA-LDL specific switched antibodies.

**Figure 1.**
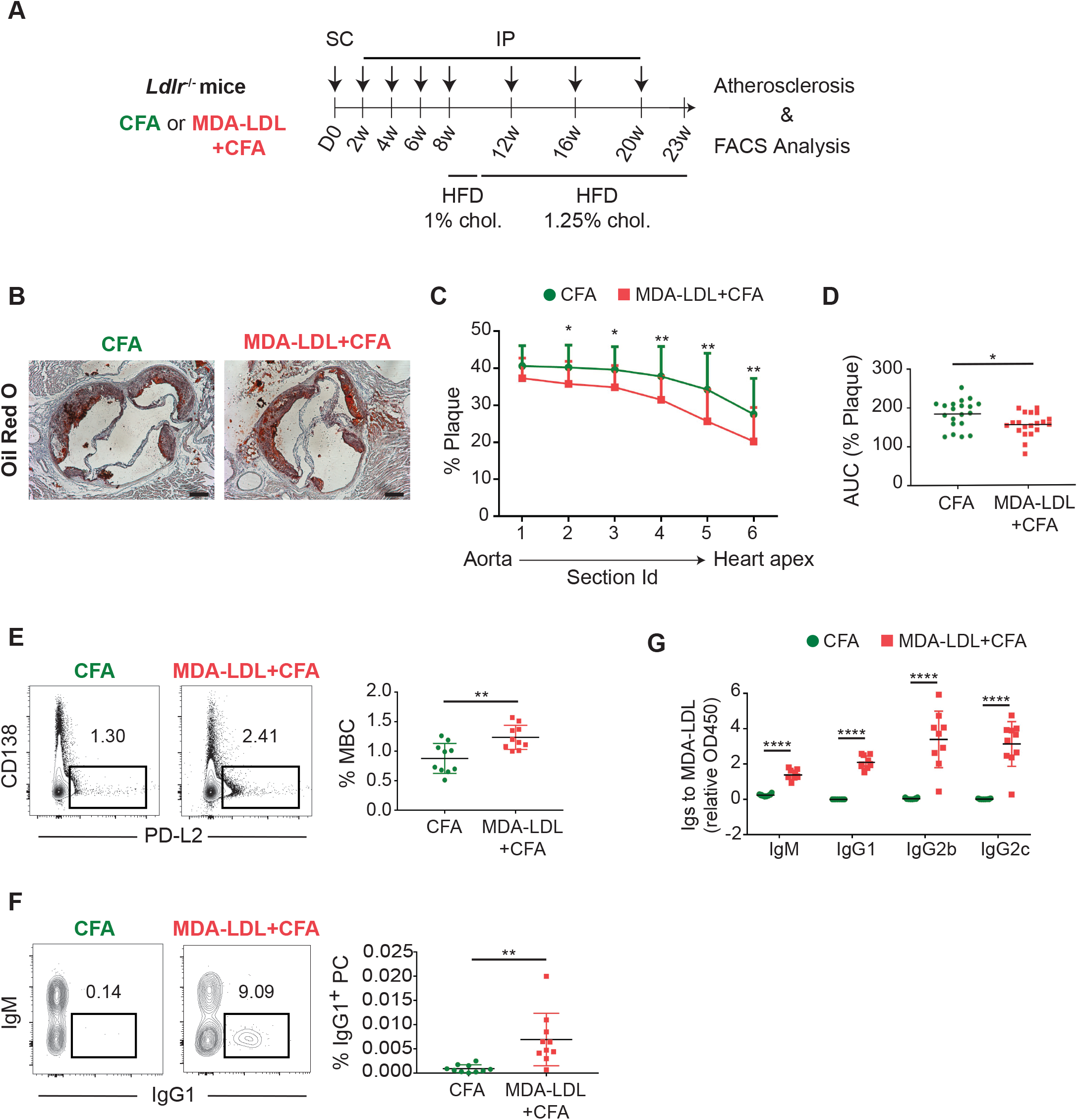
MDA-LDL is athero-protective and promotes a specific humoral response. **(A)** Experimental design. *Ldlr*^-/-^ mice were immunized subcutaneously (SC) with MDA-LDL in CFA or with CFA alone (D0). Four booster immunizations were done given intraperitoneally (IP) with MDA-LDL in IFA or IFA alone every two weeks (weeks 2, 4, 6, 8), followed by three additional immunizations once a month four weeks later (week 12, 16, 20). Mice were fed with 1% cholesterol HFD and with 1.25% cholesterol HFD from weeks 8-10 and 10-23, respectively. **(B)** Representative Oil red O/Hematoxylin staining and **(C)** quantification of the proportion of atheroma plaque in serial 80 μm-spaced aortic sinus cryosections from MDA-LDL+CFA and CFA immunized mice. **(D)** Area under the curve (AUC) plot of atheroma plaque quantification. The scale bars in images represents 200 μm. Representative flow cytometry plots and quantification of **(E)** splenic MBCs (B220^+^PD-L2^+^) and **(F)** IgG1^+^ bone marrow PCs (B220^-^CD138^+^IgG1^+^) from MDA-LDL+CFA and CFA mice. Population percentages in FACS graphs indicate the frequency of live cells. **(G)** Quantification by ELISA of MDA-LDL specific IgM, IgG1, IgG2b and IgG2c levels in serum from *Ldlr*^-/-^ mice 23 weeks after primary immunization with MDA-LDL+CFA or CFA. Data analyzed by unpaired t and two-way ANOVA statistical tests. *P≤0.05, **P<0.01 and ****P<0.0001.

To dissect the B cell immune response to MDA-LDL in the absence of atherosclerosis-dependent variables, we performed immunization experiments in *Aicda*-Cre^+/ki^ ^54,55^ *R26tdTom*^+/ki^ mice ^56^, where B cells that have been activated for Activation Induced Deaminase (AID) expression become irreversibly tdTom^+^ (Figure S2). AID is the enzyme that initiates SHM and CSR; thus, its expression is a good proxy to identify cells that have had GC experience^27,57–59^ *Aicda*-Cre^+/ki^ *R26tdTom*^+/ki^ mice were immunized with MDA-LDL+CFA or with CFA alone as shown in Figure 2A. We detected an increase in tdTom^+^GL7^+^ GC B cells and tdTom^+^PDL-2^+^ MBCs in lymph nodes and spleen of MDA-LDL+CFA immunized mice (Figure 2B-D). In addition, PCs accumulated over time in the bone marrow and were significantly increased 2 weeks after the third immunization (Figure 2E-F). This response was accompanied by an increase in both unswitched and switched anti-MDA-LDL antibody titers in MDA-LDL immunized mice, as measured by enzyme-linked immunosorbent assay (ELISA) (Figure 3A-D).

**Figure 2.**
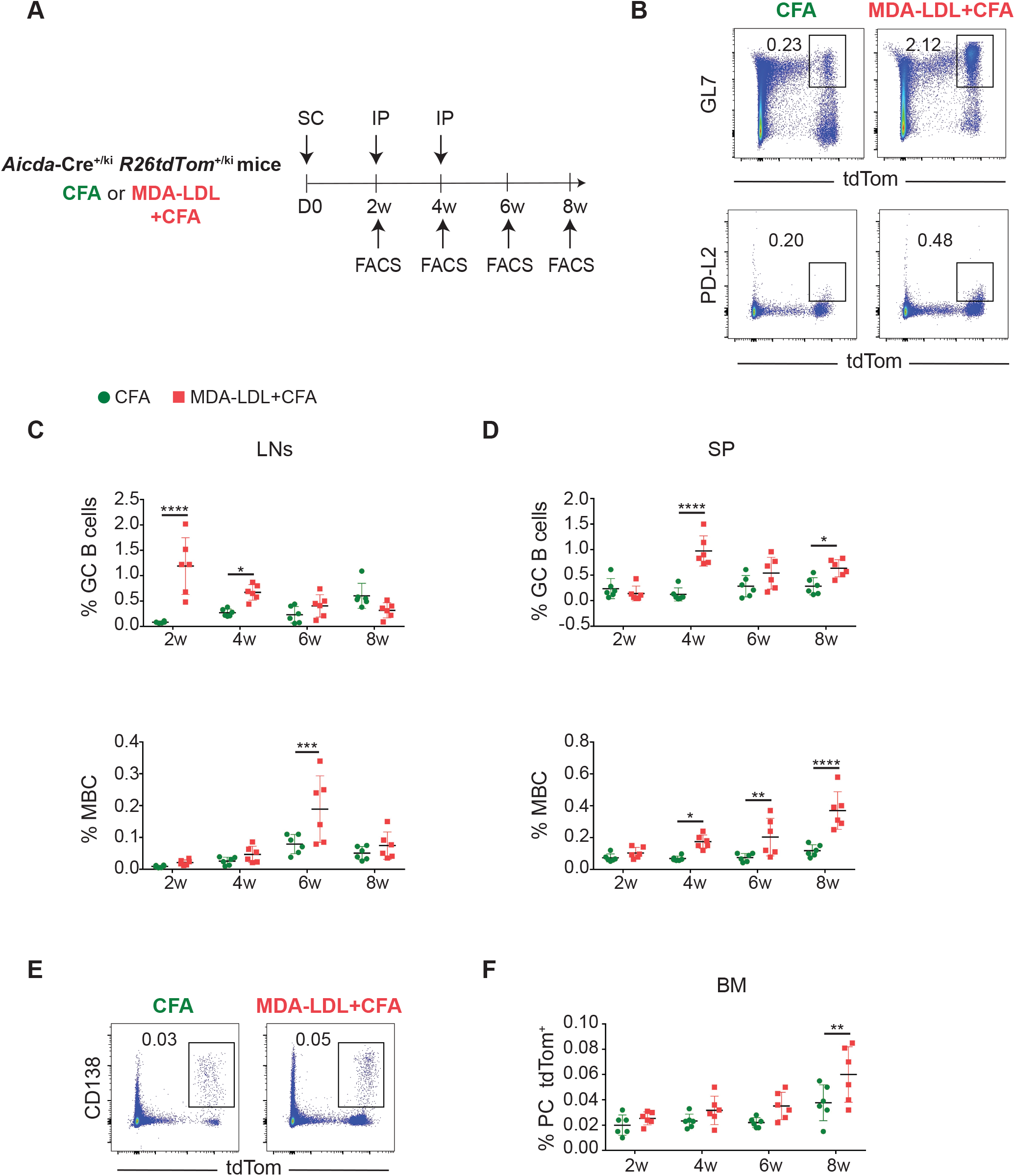
MDA-LDL immunization triggers a GC B cell response in non-atherogenic mice. **(A)** Experimental design. Mice were immunized SC with MDA-LDL in CFA or with CFA alone, followed by two booster immunizations IP with MDA-LDL in IFA or IFA alone two and four weeks later. FACS analyses were performed 2, 4, 6 and 8 weeks after the primary immunization. **(B)** Representative FACS plots of GC B cells (B220^+^GL7^+^tdTom^+^; upper panel) and MBCs (B220^+^PD-L2^+^tdTom^+^; bottom panel) 2 and 6 weeks after primary immunization, respectively and quantification of these populations in **(C)** lymph nodes (LNs) and **(D)** spleen (SP). **(E)** Representative FACS plots of bone marrow (BM) tdTom^+^ PCs (B220^-^CD138^+^tdTom^+^) 4 weeks after the primary immunization and **(F)** quantification. Population percentages in FACS graphs indicate the frequency of live cells. Data analyzed by two-way ANOVA statistical test. *P≤0.05, **P<0.01, ***P<0.001 and ****P<0.0001.

**Figure 3.**
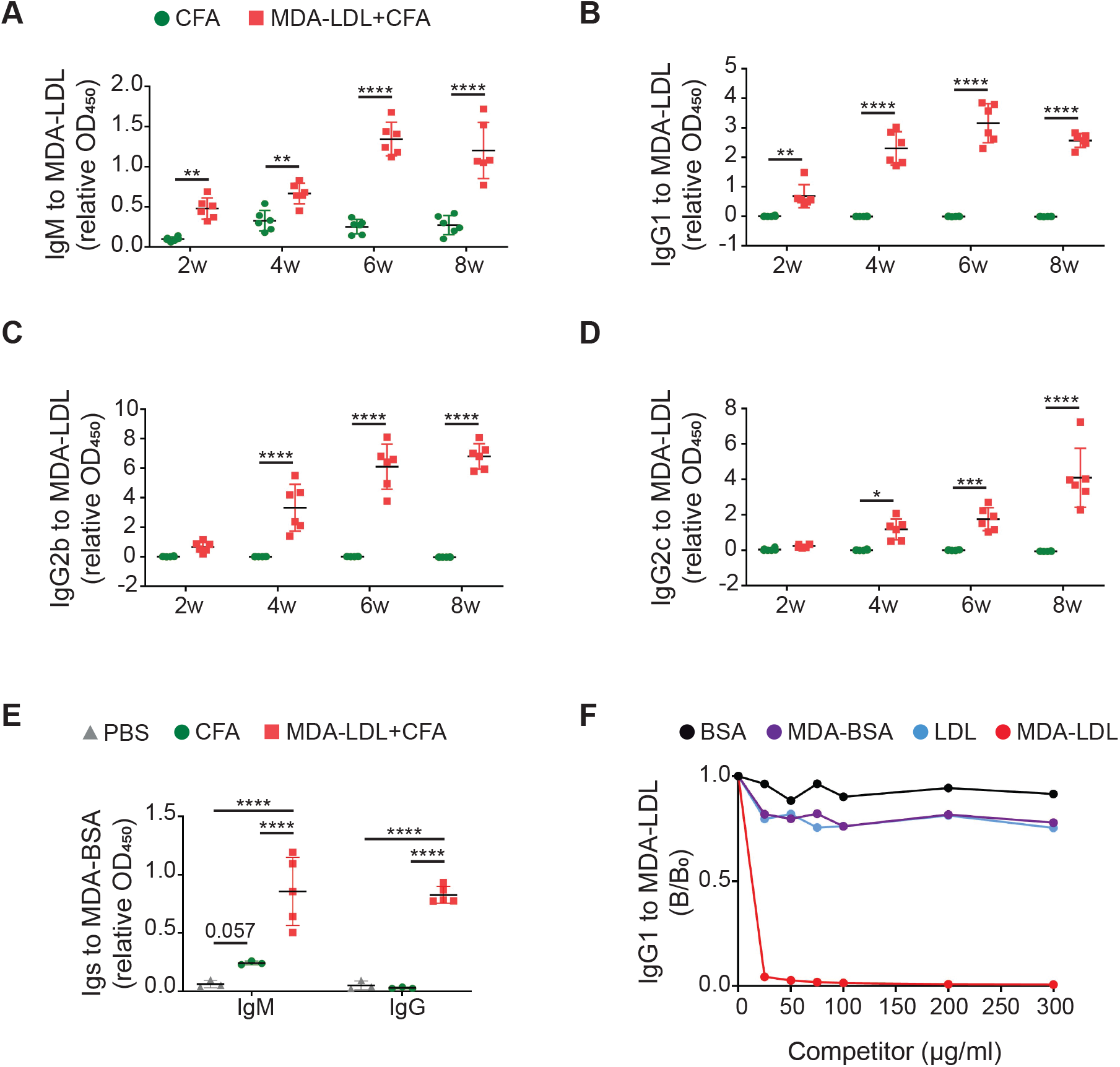
Anti-MDA-LDL antibodies recognize epitopes only presented in the MDA-modified LDL. Plasma antibody levels in *Aicda*-Cre^+/ki^ *R26tdTom*^+/ki^ mice immunized with CFA alone or MDA-LDL+CFA as determined by ELISA. Quantification of **(A)** IgM, **(B)** IgG1, **(C)** IgG2b and **(D)** IgG2c anti-MDA-LDL antibodies 2, 4, 6 and 8 weeks after primary immunization. **(E)** Quantification of IgM and IgG anti-MDA-BSA antibodies 6 weeks after primary immunization. **(F)** Competition immunoassay to measure anti-MDA-LDL IgG1 antibodies. Plasma IgG1 binding to plated MDA-LDL was quantified in the presence (B) or absence (B_0_) of increasing concentrations (0, 25, 50, 100, 150, 200 and 300 μg/ml) of the indicated competitors (BSA, MDA-BSA, LDL and MDA-LDL). Results are expressed as ratios of IgG1 binding to MDA-LDL in the presence (B) or absence (B0) of the competitor. Data analyzed by one-way and two-way ANOVA statistical tests. *P≤0.05, **P<0.01, ***P<0.001 and ****P<0.0001.

To further analyze the specificity of the antibody response to MDA-modified epitopes, we measured the antibody titers to BSA oxidized with MDA (MDA-BSA) in plasma from MDA-LDL+CFA, CFA and PBS immunized mice. We found that MDA-LDL immunization led to a dramatic increase in both IgM and IgG anti-MDA-BSA antibody titers (Figure 3E). In addition, competition immunoassays with plasma from MDA-LDL immunized mice showed that binding of plasma IgG1 antibodies to MDA-LDL was partially inhibited by pre-incubation with MDA-BSA and LDL particles, while MDA-LDL completely inhibited the binding of antibodies to MDA-LDL, indicating high specificity of the anti-MDA-LDL, in this case, IgG1 switched antibodies (Figure 3F).

Together, these results indicate that MDA-LDL immunization triggers an immune GC response that gives rise to antibodies specific for MDA-containing epitopes.

### SHM and selection in MDA-LDL immunized mice

To gain molecular insights into the antibody response to MDA-LDL we analyzed the immunoglobulin repertoire by single-cell sorting and sequencing of Ig genes in mice immunized *Aicda*-Cre^+/ki^ *R26tdTom*^+/ki^ with MDA-LDL in CFA or CFA alone. We performed single-cell sorting and sequencing of GC B cells (B220^+^ GL7^+^ tdTom^+^) and MBCs (B220^+^ GL7^-^ tdTom^+^) from lymph nodes and GC-derived PCs (B220^-^ CD138^+^ tdTom^+^) from bone marrow (Figure 4A, S3A). We obtained 597 heavy chain (IgH) and light chain (IgL) paired sequences from three mice immunized with MDA-LDL and two mice immunized with CFA alone (Table S2, Figure 4B, Figure S3B). We found minor differences in the VJ usage between the MDA-LDL+CFA and the CFA repertoires (Figure S4A-F), with VH2 family being more represented in MDA-LDL+CFA mice, and no differences in CDR3 length (Figure S4G-H). Conversely, VH5 and VH10 families were under-represented in the MDA-LDL+CFA repertoire (Figure S4A).

**Figure 4.**
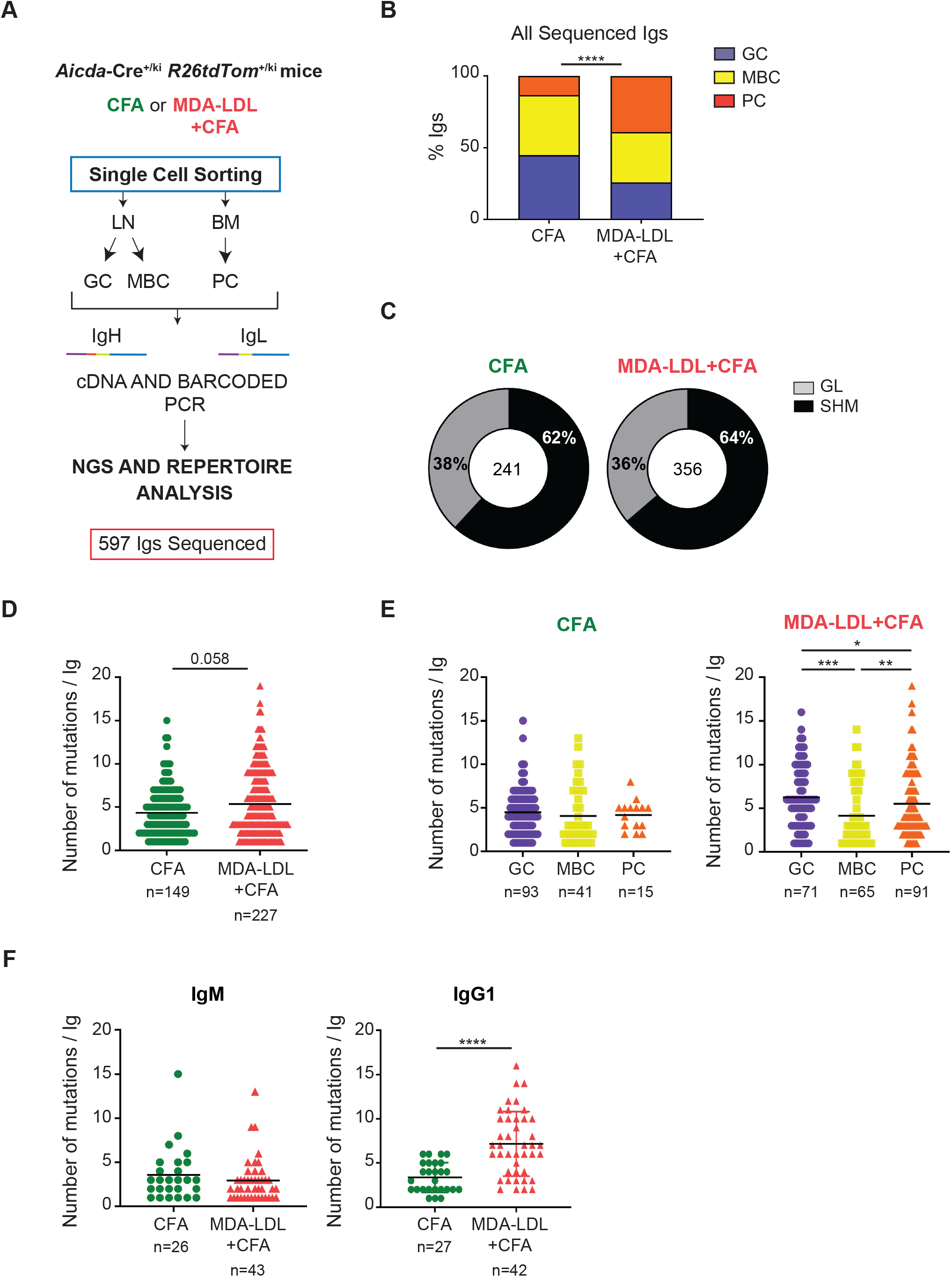
MDA-LDL immunization induces highly mutated IgG1 antibodies. **(A)** GC B cells and MBCs from lymph nodes and GC-derived PCs (PC tdTom+) from bone marrow of CFA (n=2) and MDA-LDL+CFA (n=3) immunized mice (3 times every two weeks) were isolated 2 weeks after last boost by flow cytometric single-cell sorting. IgH and IgL genes were amplified and sequenced. Paired Igh and Igk/Igl gene sequences from a total of 597 Igs were obtained and analyzed. **(B)** Population distribution of all Igs sequenced within each group. **(C)** Proportion of mutated (SHM) and germ line (GL) Igs in CFA and MDA-LDL+CFA repertoires. Number of IgH somatic mutations (nucleotide exchanges) within mutated Igs in **(D)** all cells, **(E)** by cell population and **(F)** in IgM and IgG1 B cells from each group. Data analyzed by non-parametric Mann-Whitney statistical test and Chi-Square test. *P≤0.05, **P<0.01, ***P<0.001 and ****P<0.0001.

Regarding SHM, we did not observe significant differences in the proportion of mutated Igs or in their overall mutation frequency between the CFA and the MDA-LDL+CFA group (Figure 4C-D). However, while GC, MBC and PC subsets showed similar SHM frequencies in the CFA group, in MDA-LDL immunized mice, Igs from GC B cells were the most mutated, followed by the Igs from PCs, and with Igs from MBCs harbored the fewest mutations (Figure 4E). Moreover, only IgG1+ cells, but no other isotypes, showed a significant increase of SHM in MDA-LDL immunized mice compared to control mice (Figure 4F, S5A-E) and MDA-LDL+CFA immunized mice had a larger proportion of GC IgG1+ cells (Figure S5F-H). Together, these data indicate that MDA-LDL immunization triggers an immune response that involves a GC reaction.

To focus on Igs relevant to the MDA-LDL response, we identified those expressed by clonally expanded B cells. Thus, we defined “B cell clones” as clusters of three or more Igs sharing the same V and J genes and the same CDR3 length in both IgH and IgL. We identified 25 B cell clones, 12 of which were exclusively comprised by Igs from CFA group (1-12), 12 by Igs from MDA-LDL+CFA group (13-24) (from now on, exclusive clones) and 1 clone containing Igs from both groups (25) (mixed clone) (Table S3 and Figure 5A).

**Figure 5.**
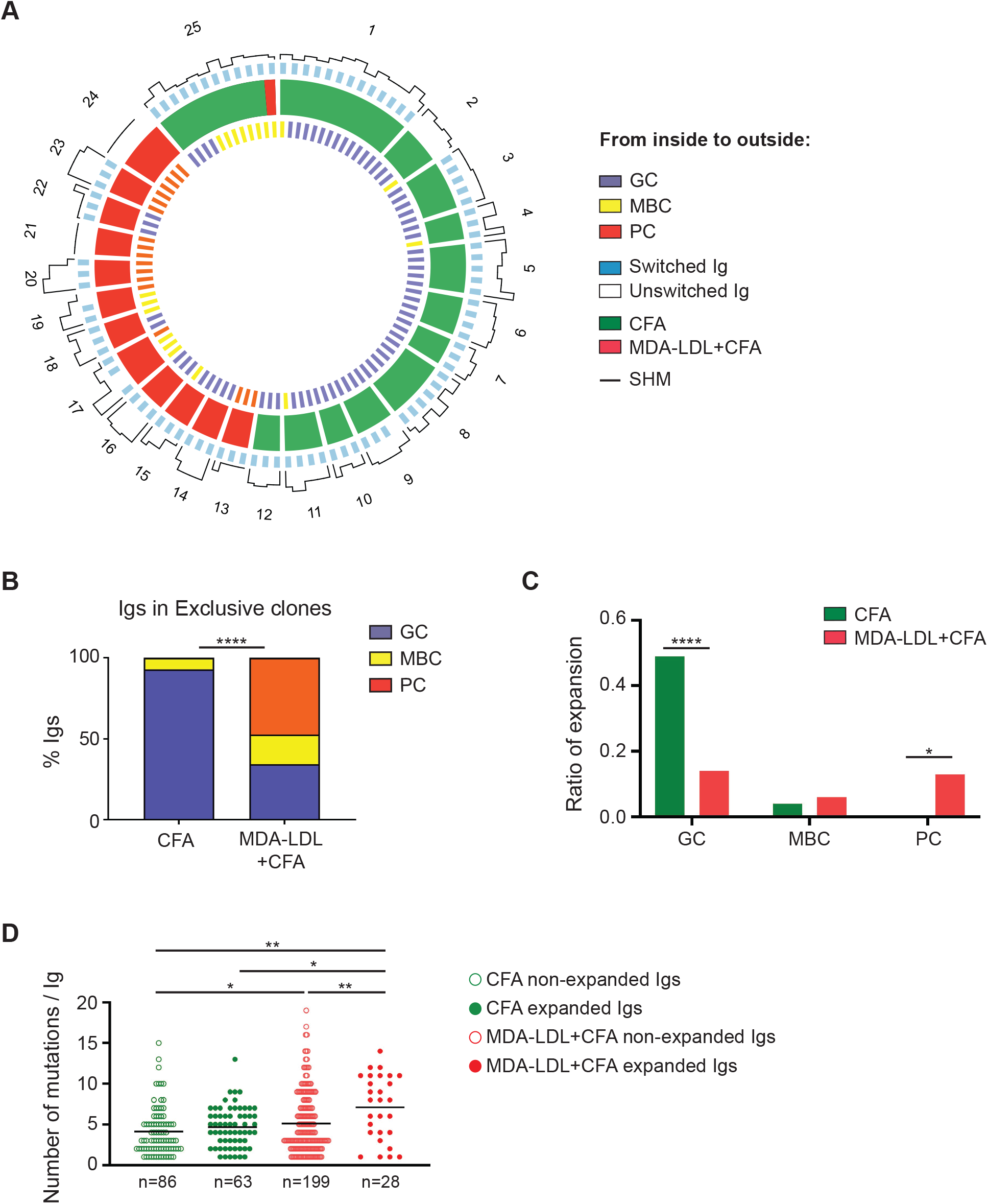
Igs from clonally expanded B cell clones are highly mutated after MDA-LDL+CFA immunization. **(A)** Circos plots illustrate the molecular features of expanded Igs. Igs from clonally expanded cell clones are represented as boxes fill in green (CFA exclusive clones), red (MDA-LDL+CFA exclusive clones) and both colors (shared clones). Clone number is shown in the outer layer. Cell population origin, CSR and number of Ig mutations (SHM) (nucleotide exchanges) is indicated for each B cell. **(B)** Population distribution of Igs present in exclusive clones from each group. **(C)** Ratio of expansion (expanded cells/all cells) of each cell population. **(D)** Number of somatic mutations (nucleotide exchanges) per mutated nonexpanded Ig or expanded Ig in each group. Data were analyzed by non-parametric Kruskal-Wallis statistical, Fisher’s exact and Chi-Square tests. *P≤0.05, **P<0.01 and ****P<0.0001.

We found that exclusive CFA clones were disproportionally comprised by Igs expressed by GC B cells while MDA-LDL+CFA clones were composed by Igs from GC, PC and MBC populations (Figure 5B) and showed a greater enrichment in PC (Figure 5C). Finally, while in CFA mice we did not observe differences in mutation frequency between expanded and non-expanded Igs, in MDA-LDL+CFA mice expanded Igs had higher mutational load than non-expanded Igs (Figure 5D). This result shows that B cells expanded in response to MDA-LDL express heavily mutated Igs, indicating that they have undergone affinity maturation.

### GC-derived plasma cells are athero-protective

To address the role of GC-derived antibodies in atherosclerosis, we transferred bone marrow cells from *Prdm1*^fl/fl^ *Aicda*-Cre^+/ki^ mice into lethally irradiated *Ldlr*^-/-^ mice (from now on, *Prdm1*^fl/fl^ *Aicda*-Cre^+/ki^ *Ldlr*^-/-^ chimeras). *Prdm1* codes for Blimp1, the master transcriptional regulator of PCs^60^; thus, *Prdm1*^fl/fl^ *Aicda*-Cre^+/ki^ *Ldlr*^-/-^ chimeras will be devoid of PCs and antibody secretion derived of cells that have expressed AID. *Ldlr*^-/-^ mice transferred with *Prdm1*^+/+^ *Aicda*-Cre^+/ki^ bone marrow (from now on, *Prdm1*^+/+^ *Aicda*-Cre^+/ki^ *Ldlr*^-/-^ chimeras) were used as controls (Figure 6A). Following bone marrow reconstitution, chimeras were fed with high fat diet (HFD) for nine weeks. We found no significant differences in weight gain (Figure S6A) or plasma lipid profile (Table S4) between the two groups.

**Figure 6.**
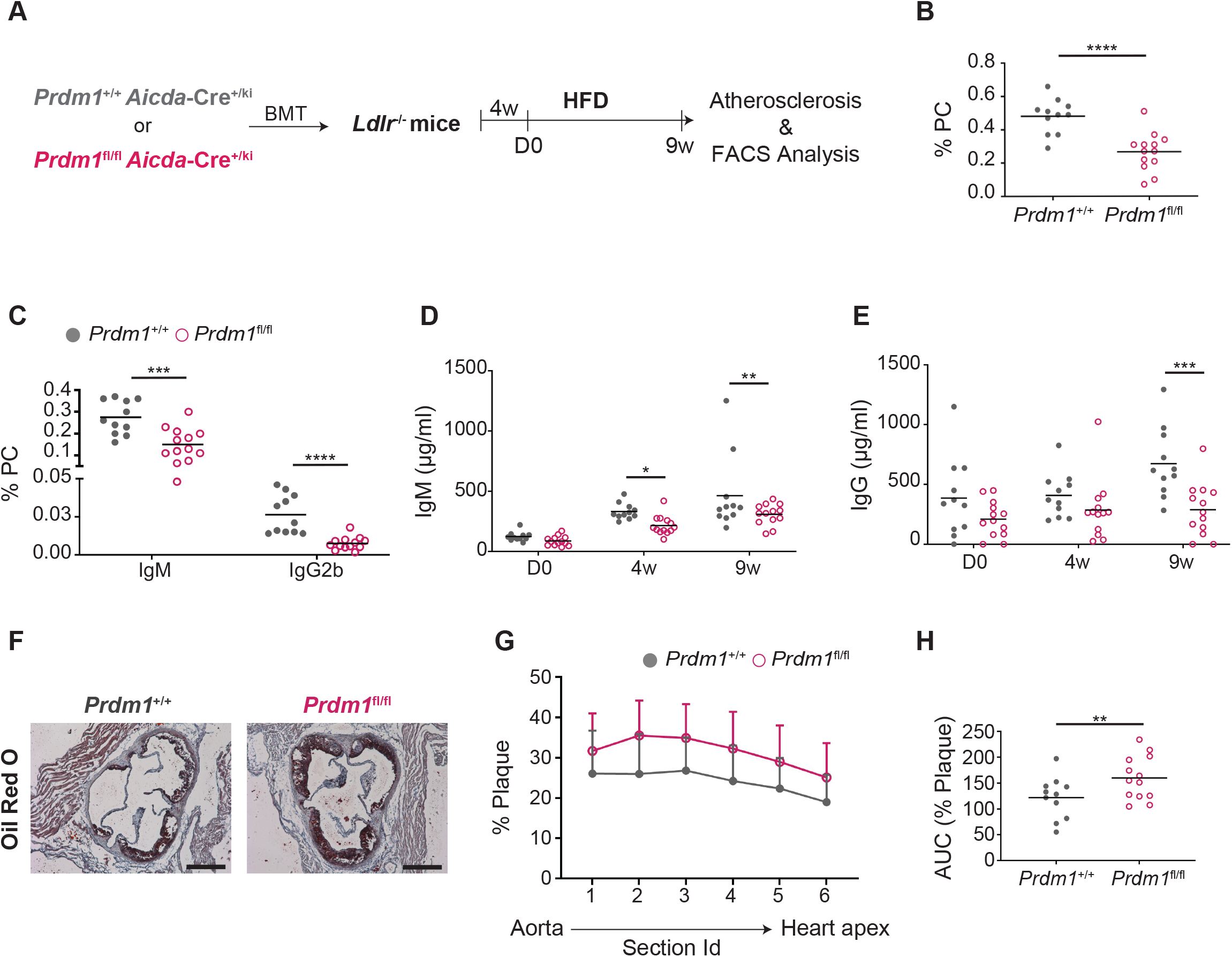
*Prdm1*^fl/fl^ *Aicda*-Cre^+/ki^ *Ldlr*^-/-^ chimeras show increased atherosclerosis. **(A)** Experimental design. Bone marrow from *Prdm1*^+/+^ *Aicda*-Cre^+/ki^ or *Prdm1*^fl/fl^ *Aicda*-Cre^+/ki^ mice was isolated and intravenously (i.v.) transferred to previously irradiated *Ldlr*^-/-^ mice. 4 weeks after bone marrow transfer, mice were fed with 0.2% cholesterol HFD for 9 weeks. Quantification of **(B)** PC percentage (B220^-/low^ CD138^+^) and **(C)** IgM positive (B220^-/low^ CD138^+^ IgM^+^) and IgG2b positive (B220^-/low^ CD138^+^ IgG2b^+^) PC in spleen. Quantification of total **(D)** IgM and **(E)** IgG antibody titers in plasma at day 0, 4 and 9 weeks of HFD as measured by ELISA. **(F)** Representative Oil red O/Hematoxylin stains and **(G)** quantification of the proportion of atheroma plaque in serial 80 μm-spaced aortic sinus cryosections from *Prdm1*^+/+^ and *Prdm1*^fl/fl^ *Aicda*-Cre^+/ki^ *Ldlr*^-/-^ chimeras. **(H)** AUC plot of atherosclerosis quantification. The cale bars in images represents 200 μm. Data analyzed by unpaired t statistical test and twoway ANOVA statistical test. *P≤0.05, **P<0.01, ***P<0.001 and ****P<0.0001.

Expectedly, the proportion of CD138^+^ PC was reduced in the spleens of *Prdm1*^fl/fl^ *Aicda*-Cre^+/ki^ *Ldlr*^-/-^ chimeras compared to control mice (Figure 6B). This reduction was more severe in IgG2b isotype-switched PC (Figure 6C); the CD138^+^IgM^+^ PCs remaining in *Prdm1*^fl/fl^ *Aicda*-Cre^+/ki^ *Ldlr*^-/-^ chimeras probably represent PC derived from extrafollicular B cells. Likewise, plasma IgM and IgG antibody titers were reduced in Prdm1-deficient chimeras (Figure 6D-E). These results indicate that a large proportion of PCs, including those derived from GC B cells, are depleted in our *Prdm1*^fl/fl^ *Aicda*-Cre^+/ki^ *Ldlr*^-/-^ model.

To assess the contribution of GC-derived PC on atherosclerosis, we evaluated atherosclerosis progression in *Prdm1*^fl/fl^ *Aicda*-Cre^+/ki^ *Ldlr*^-/-^ and *Prdm1*^+/+^ *Aicda*-Cre^+/ki^ *Ldlr*^-/-^ chimeras. Atherosclerosis quantification in aortic sinus revealed a significant increase in atherosclerosis in Prdm1-deficient chimeras compared to controls (Figure 6F-H). Atherosclerosis acceleration did not associate with obvious differences in atheroma plaque composition as measured by staining of collagen content (Figure S6B-C), as well as macrophages and smooth muscle cells (SMCs) immunofluorescence (Figure S6D-F). We conclude that GC-derived PCs and/or antibodies protect from atherosclerosis development.

### GC-derived antibodies contribute to MDA-LDL immunization mediated athero-protection

To assess the role of GC-derived antibodies in the MDA-LDL mediated athero-protection, we immunized Prdm1-deficient pro-atherogenic chimeras with MDA-LDL. Briefly, we transferred bone marrow from *Prdm1*^fl/fl^ *Aicda*-Cre^+/ki^ or *Prdm1*^+/+^ *Aicda*-Cre^+/ki^ mice into *Ldlr*^-/-^ mice, and reconstituted chimeras were immunized with MDA-LDL+CFA, CFA or PBS and fed with HFD as indicated in Figure 7A. Chimeras from all 6 groups showed similar weight gain (Figure S7). Plasma levels of total cholesterol, free cholesterol, LDL cholesterol and triglyceride pro-atherogenic indexes^61^, but not high-density lipoprotein (HDL) cholesterol, tended to be reduced in CFA and MDA-LDL+CFA immunized chimeras (Figure S8). Reduction in pro-atherogenic lipid levels were observed both in Prdm1-proficient and -deficient chimeras, indicating that modulation of plasma lipid profile seems to be independent of GC-derived Abs.

**Figure 7.**
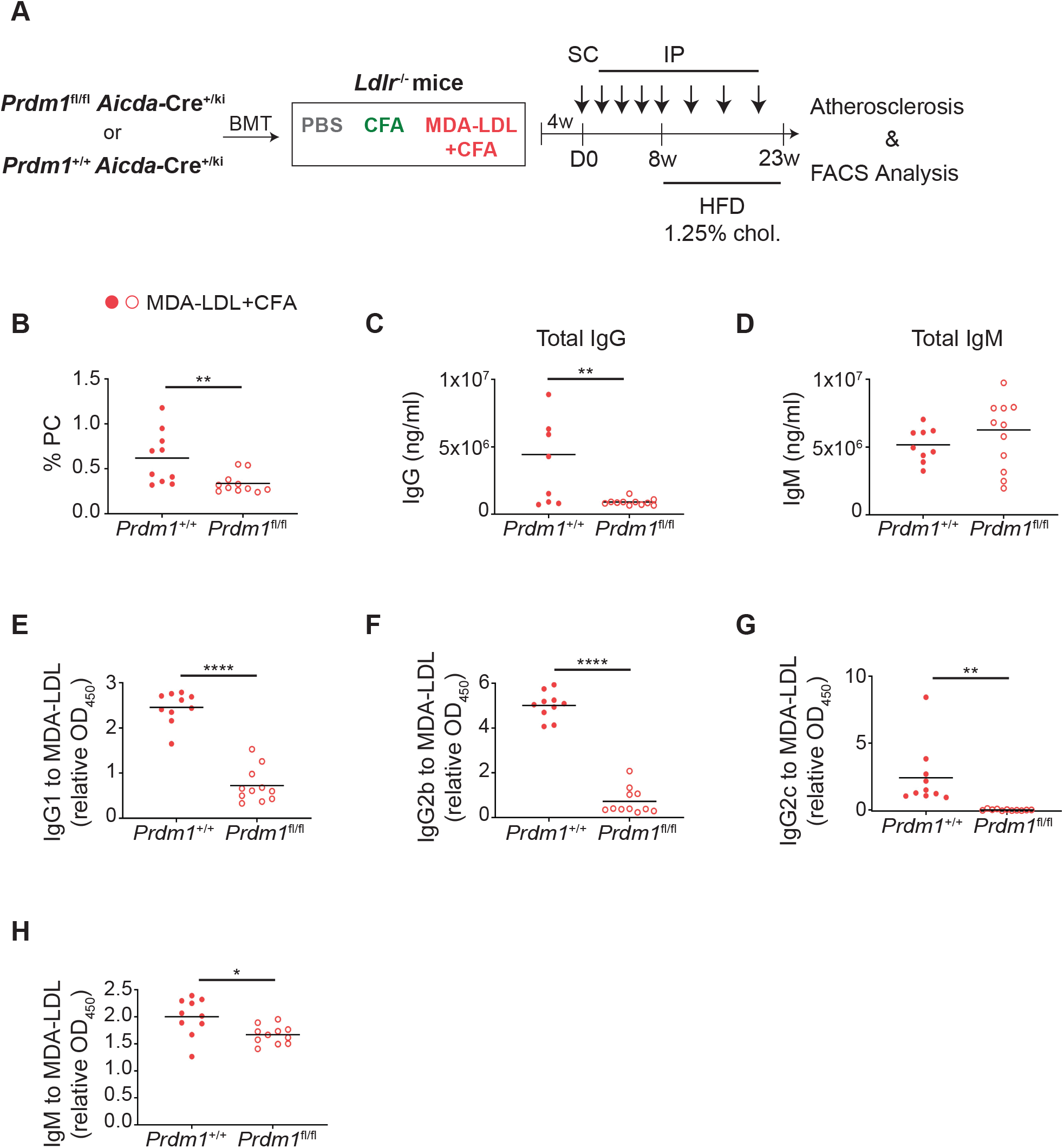
Low MDA-LDL-specific antibody responses in the absence of GC-derived PCs. **(A)** Experimental design. Bone marrow from *Prdm1*^+/+^ *Aicda*-Cre^+/ki^ or *Prdm1*^fl/fl^ *Aicda*-Cre^+/ki^ mice was i.v. transferred to into irradiated *Ldlr*^-/-^ recipients. 4 weeks later, mice were immunized with MDA-LDL+CFA, CFA alone or PBS and fed with HFD with 1.25% cholesterol for 15 weeks. 23 weeks after the primary immunization, mice were sacrificed for analysis. **(B)** Quantification of spleen PCs (B220^-/low^ CD138^+^) from MDA-LDL+CFA immunized *Prdm1*^+/+^ and *Prdm1*^fl/fl^ *Aicda*-Cre^+/ki^ *Ldlr*^-/-^ chimeras by FACS. ELISA-based quantification of total **(C)** IgG and **(D)** IgM antibody levels in plasma from *Prdm1*^+/+^ and *Prdm1*^fl/fl^ *Aicda*-Cre^+/ki^ *Ldlr*^-/-^ chimeras immunized with MDA-LDL+CFA. MDA-LDL-specific **(E)** IgG1, **(F)** IgG2b, **(G)** IgG2c and **(H)** IgM antibody levels in *Prdm1*^+/+^ and *Prdm1*^fl/fl^ *Aicda*-Cre^+/ki^ *Ldlr*^-/-^ chimeras immunized with MDA-LDL+CFA measured by ELISA. Data analyzed by unpaired t statistical test. *P≤0.05, **P<0.01 and ****P<0.0001.

Expectedly, we found that the PC population was reduced in the spleen of MDA-LDL+CFA immunized *Prdm1*^fl/fl^ *Aicda*-Cre^+/ki^ *Ldlr*^-/-^ compared to immunized control chimeras (Figure 7B). Plasma titers of IgG antibodies were also reduced in MDA-LDL immunized *Prdm1*^fl/fl^ *Aicda*-Cre^+/ki^ *Ldlr*^-/-^ chimeras (Figure 7C), while total IgM titers were not significantly changed (Figure 7D). Likewise, Prdm1-deficient pro-atherogenic chimeras showed a sharp decrease in anti-MDA-LDL IgG1, IgG2b and IgG2c Ab titers (Figure 7E-G) and a mild reduction of anti-MDA-LDL IgM antibody titers (Figure 7H). These results indicate that PCs derived from GCs play a critical role in the antibody response triggered by MDA-LDL immunization in atherosclerotic mice.

To test the role of GC-derived Abs in atherosclerosis, we then measured atherosclerotic extent in aortic sinus of *Prdm1*^+/+^ *Aicda*-Cre^+/ki^ *Ldlr*^-/-^ and *Prdm1*^fl/fl^ *Aicda*-Cre^+/ki^ *Ldlr*^-/-^ chimeras immunized with MDA-LDL in CFA or CFA alone (experiment depicted in Figure 7A). We included a group of mice injected with PBS to normalize for reconstitution variability between groups (Figure 8A-B). We found that both *Prdm1*^fl/fl^ *Aicda*-Cre^+/ki^ *Ldlr*^-/-^ and *Prdm1*^fl/fl^ *Aicda*-Cre^+/ki^ *Ldlr*^-/-^ chimeras treated with CFA developed atherosclerotic plaques of similar size (Figure 8C), indicating that CFA atheroprotection is independent of GC-derived antibodies. In contrast, upon MDA-LDL immunization, *Prdm1*^fl/fl^ *Aicda*-Cre^+/ki^ *Ldlr*^-/-^ chimeras had larger aortic plaque lesions than did Prdm1-proficient chimeras (Figure 8D). Therefore, these results indicate that PCs and antibodies derived from GC-experienced B cells drive MDA-LDL mediated athero-protection.

**Figure 8.**
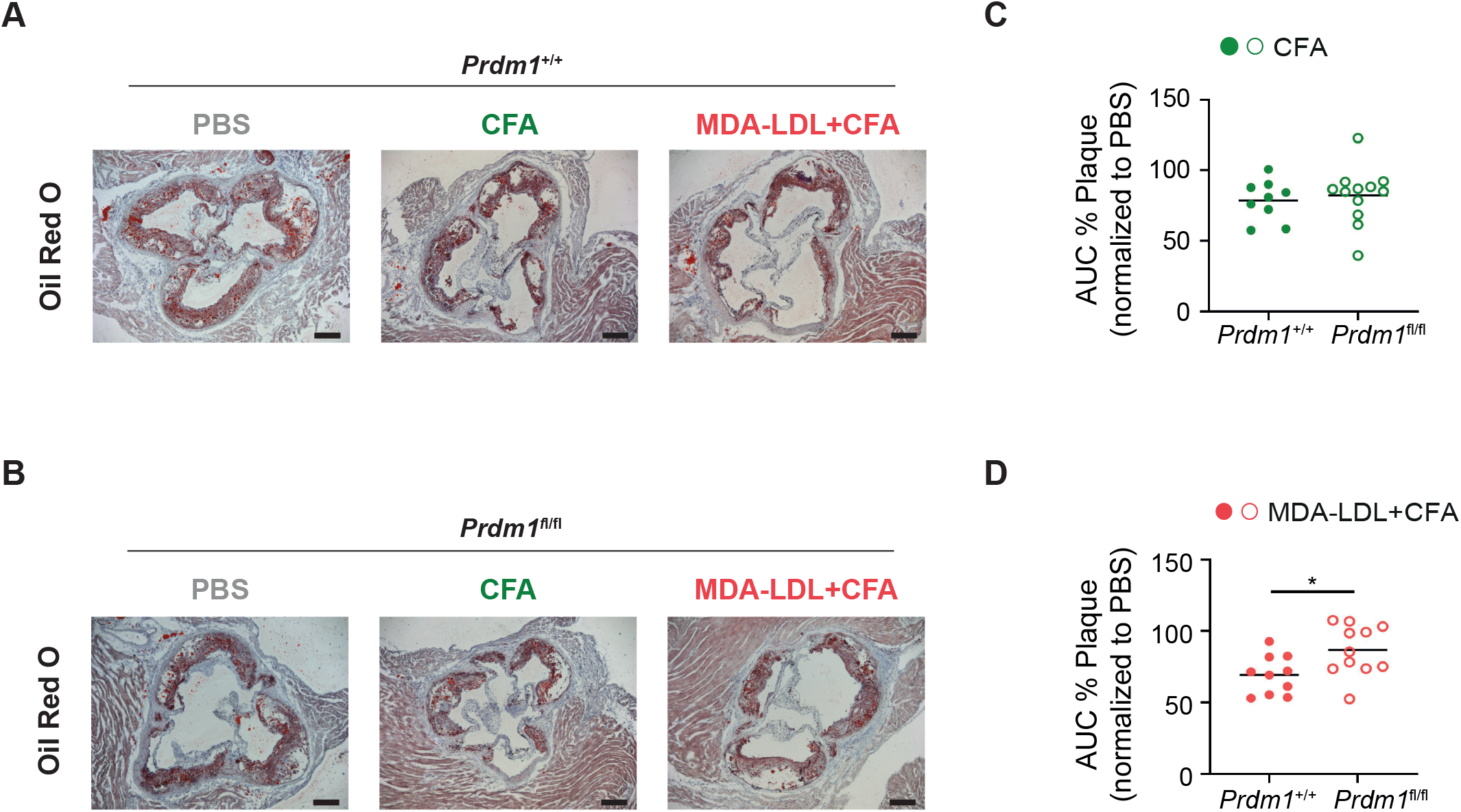
GC-derived PCs and antibodies are required for MDA-LDL-immunization induced athero-protection. Representative Oil Red-O-stained aortic sinus cryosections of **(A)** *Prdm1*^+/+^ *Aicda*-Cre^+/ki^ *Ldlr*^-/-^ chimeras and **(B)** *Prdm1*^fl/fl^ *Aicda*-Cre^+/ki^ *Ldlr*^-/-^ chimeras immunized with PBS, CFA, or MDA-LDL+CFA. AUC values of the atherosclerosis quantifications in five serial 80 μm-spaced aortic sinus cryosections from *Prdm1*^+/+^ *Aicda*-Cre^+/ki^ *Ldlr*^-/-^ and *Prdm1*^fl/fl^ *Aicda*-Cre^+/ki^ *Ldlr*^-/-^ chimeras immunized with **(C)** CFA or **(D)** MDA-LDL+CFA normalized to the mean AUC value of PBS immunized chimeras of same genotype. Scale bars in images represents 200 μm. Data analyzed by unpaired t statistical test. *P<0.05.

## DISCUSSION

Here we have shown that MDA-LDL immunization of proatherogenic *Ldlr*^-/-^ mice triggers an antibody immune response which entails the generation of GCs, switched antibodies, MBCs and PCs. Our experiments in *Aicda*-Cre^+/ki^ *R26tdTom*^+/ki^ mice have allowed for the first time to trace activated B cells after MDA-LDL immunization, and has confirmed this GC response in a non-atherogenic background, thus eliminating the confounding effects that the atherosclerosis immune response might have. Interestingly, we detected antibodies specific of the MDA moiety in the context of the LDL particle. Thus, although native LDL by itself can raise an immune response^63^ our data support that MDA-modified LDL is indeed immunogenic, in agreement with previous data ^52^. Of note, our results were obtained by comparing MDA-LDL in CFA with mice immunized with CFA alone, which is also known to promote athero-protection ^62^. Interestingly, MDA-LDL promotes an increase in the load of SHM, particularly in expanded clones of B cells, strongly suggesting that affinity maturation is in place in MDA-LDL immunized mice and therefore, that MDA-LDL, but not CFA alone, triggers a *bona fide* GC response, much like other antigenic triggers used for vaccination strategies ^64,65^. In this regard, it is worth mentioning that the GLACIER trial (Goal of Oxidized LDL and Activated Macrophage Inhibition by Exposure to a Recombinant Antibody) was approved to assess the atheroprotective properties of passive immunization with the monoclonal anti-MDA MLDL1278A antibody. The results did not reveal atheroprotective effects, at least in the cohort of subjects with stable CVD analyzed in the trial^6,66^. However, it is tempting to suggest that additional antibodies, such as the expanded clones identified in the present study, can also prove valuable candidates in future clinical trials.

In our study we have addressed the general contribution of GC-derived antibodies using *Prdm1*^fl/fl^ *Aicda*-Cre^+/ki^ mice. In this model, cells that have activated AID expression are depleted of *Prdm1*. We found that *Prdm1*^fl/fl^ *Aicda*-Cre^+/ki^ *Ldlr*^-/-^ chimeras showed increased atherosclerosis, indicating an athero-protective effect of GC-derived PCs and/or antibodies. Two *Prdm1* models have been previously reported in the context of atherosclerosis with diverse results, which may be explained by the different experimental strategies for *Prdm1* deletion. In one study, *Prdm1* deletion was driven by a *Cd23*-Cre, thus driving effective depletion of Blimp1 from all follicular-derived B cells ^26^. In a second study, *Prdm1* deletion was driven by the *Cd19*-Cre allele, leading to Blimp1 deletion in the vast majority of B cells^27^. Although the onset of AID expression shortly precedes B cell entry into the GC, our approach allows a more restricted depletion of GCs derived PCs than previous reports. This is further indicated by the persistance of a considerable fraction of IgM+ PCs in *Prdm1*^fl/fl^ *Aicda*-Cre^+/ki^ mice. Therefore, it is possible that antibody secreting cells depleted in the *Cd19*-Cre or *Cd23*-Cre models -but not in *Prdm1*^fl/fl^ *Aicda*-Cre^+/ki^ mice-have pro-atherogenic properties. Interestingly, abolishing antibody secretion by deletion of *Xbp-1* with *mb1*-Cre, which is expressed from early B cell precursors, resulted in increased atherosclerosis, as did depletion of T cells with anti-CD4 antibodies^28^. These results suggest that the endogenous T cell-dependent response can be athero-protective^28^, in agreement with our results. In addition, the differential impact on atherosclerosis observed in these models may be the consequence of a subtle functional balance between antibodies with different antigen specificities, some of which can have athero-protective properties, while others can aggravate the disease. For instance, anti-ALDH4A1 antibodies are athero-protective^34^, while anti-hsp60 antibodies aggravate atherosclerosis^67^.

In this regard, we have assessed the role of MDA-LDL specific PC and/or antibodies on atherosclerosis development by MDA-LDL immunization of proatherogenic mice under deletion of Blimp1 in GC-derived PCs (*Prdm1*^fl/fl^ *Aicda*-Cre^+/ki^ *Ldlr*^-/-^ chimeras). Depletion of GC-derived PCs dramatically decreased the titers of IgG antibodies in MDA-LDL immunized mice, and MDA-LDL-specific IgG antibodies were barely detectable. In contrast, titers of total IgM antibodies in MDA-LDL immunized *Prdm1*^fl/fl^ *Aicda*-Cre^+/ki^ *Ldlr*^-/-^ chimeras remain unchanged compared to PC proficient mice, and anti-MDA-LDL-specific IgM antibodies were only mildly reduced. Therefore, our approach allows a precise study of antibody contribution to MDA-LDL protection because it successfully abolishes the generation of GC-derived, switched antibodies while retaining a substantial proportion of IgM antibodies, presumably derived from extrafollicular B cell activation. Our results show that MDA-LDL immunization failed to be atheroprotective Prdm1-deficient chimeras. Thus, our study indicates that MDA-LDL immunization gives rise to athero-protective antibodies of GC origin.

Athero-protection by MDA-LDL immunization was first described more than two decades ago^44^, opening new therapeutic perspectives based inthrough immunomodulation strategies, and most specifically via vaccination strategies using MDA-LDL. However, our limited knowledge on the nature of the immune response to MDA-LDL and its impact on the atherosclerosis process has hindered further progress of this potential therapeutic avenue. Here, we have shown that immunization with MDA-LDL promotes a *bona fide* GC response and generates MBCs, PCs and highly mutated anti-MDA-LDL antibodies, which are important drivers of MDA-LDL-derived athero-protection. Our data thus indicate that MDA-LDL immunization can warrant a high affinity, long-lived antibody immune response, possibly comparable to the response elicited by antigens used in vaccination protocols.

## METHODS

### Mice

*Ldlr*^-/-^ mice were originally obtained from Jackson Laboratories (002207) (Ishibashi et al., 1993). *Aicda*-Cre^+/ki^ *R26tdTom*^+/ki^ mice were generated by crossing *Aicda*-Cre^+/ki^ mice obtained from Jackson Laboratories (007770) ^54^ with *R26tdTomato*^+/ki^ mice acquired from Jackson Laboratories (007914)^56^. *Prdm1*^fl/fl^ mice were acquired from Jackson Laboratories (008100)^68^ and crossed with *Aicda*-Cre^+/ki^ mice to generate *Prdm1*^*f*l/fl^ *Aicda*-Cre^+/ki^ and control *Prdm1*^+/+^ *Aicda*-Cre^+/ki^ mice. *Ldlr*^-/-^ males were used for atherosclerosis experiments (Figure 1, and host *Ldlr*^-/-^ for BMT experiments in Figures 7 and 8) in order to reduce the variability of atherosclerosis progression between males and females. Both males and females were used in the rest of the experiments. For MDA-LDL athero-protection experiments, mice were fed with HFD containing 21% of crude fat and 1% cholesterol and/or with HFD containing 21% of crude fat and 1.25% cholesterol (EF D12079 mod. 1% chol and EF D12079 mod. 1.25% chol, Ssniff Spezialdiäten, respectively). In the rest of atherosclerosis experiments, mice were fed with a HFD containing 21% of crude fat and 0.2% cholesterol (EF D12079, Ssniff Spezialdiäten). All animals were housed at the CNIC animal facility under a 12 h dark/light cycle with food and water ad libitum. All animal procedures were approved by the CNIC Ethics Committee and the Madrid regional authorities (PROEX 377/15) and conformed to EU Directive 2010/63/EU and Recommendation 2007/526/EC regarding the protection of animals used for experimental and other scientific purposes, enforced in Spanish law under Real Decreto 1201/2005.

### Antigen preparation and immunization protocols

LDL was isolated as previously described ^69^. In brief, pooled plasma from healthy donors was mixed with different densities potassium bromide solutions and sequentially ultracentrifugated. The LDL fraction was then isolated at a density of 1.019-1.063 g/mL and dialyzed in PBS. MDA-LDL and MDA-BSA were prepared as described ^70^. Briefly, LDL or BSA (Sigma-Aldrich) were incubated with 0.5M MDA in a ratio of 100 μl MDA/mg LDL during 3 h at 37 ºC. 0.5 M MDA was freshly prepared by acid hydrolysis of malonaldehyde bis dimethylacetal (Sigma-Aldrich). After incubation, MDA-LDL and MDA-BSA were purified using PD10 desalting columns following the manufacturer’s protocol (Sigma-Aldrich, GE17-0851-01). In athero-protection experiments, 21–28-week-old *Ldlr*^-/-^ mice and 14-18-week-old *Prdm1*^+/+^ *Aicda*-Cre^+/ki^ *Ldlr*^-/-^ and *Prdm1*^fl/fl^ *Aicda*-Cre^+/ki^ *Ldlr*^-/-^ chimeras were immunized subcutaneously (footpads) with 50 μg of MDA-LDL (25 μg/footpad) emulsified in complete Freund’s Adjuvant (CFA) in a 1:1 ratio. Booster immunizations were performed 4 times every 2 weeks and 3 times more monthly, injecting 25 μg of MDA-LDL emulsified in incomplete Freund’s Adjuvant (IFA) intraperitoneally. In MDA-LDL immunization kinetics analysis, 7-10-week-old *Aicda*-Cre^+/ki^ *R26tdTom*^+/ki^ mice, were immunized as described above. Control mice were injected with PBS emulsified in CFA or IFA (1:1).

### Flow Cytometry

Single-cell suspensions were obtained from lymph nodes, spleen and bone marrow. After erythrocyte lysis, Fc receptors were blocked with anti-mouse CD16/CD32 antibodies and stained with fluorophore or biotin-conjugated anti-mouse antibodies (BD Pharmingen, Biolegend, Tonbo or eBioscience) to detect B220 (RA3-6B2), GL7 (GL-7), PD-L2 (TY25), CD138 (281-2), IgD (11-26), IgG1 (A85-1), IgG2b (RMG2b-1), IgM (IL/41). Fluorophore-conjugated streptavidin (BD) was used to detect biotin-conjugated antibodies. For live cells detection, staining with 7AAD (BD Pharmingen) or LIVE/DEAD Fixable Yellow Dead Cell Stain (Thermo Fisher) was performed. Samples were acquired on LSRFortessa or FACSCanto instruments (BD Biosciences) and analyzed with FlowJo V10.4.2 software

### Bone marrow transfer

CD45.1 *Ldlr*^-/-^ mice were lethally irradiated with two doses of 5.5 Gy. The following day, femurs and tibias from CD45.2 *Prdm1*^+/+^ *Aicda*-Cre^+/ki^ and CD45.2 *Prdm1*^fl/fl^ *Aicda*-Cre^+/ki^ mice were harvested and processed to obtain bone marrow cell suspension. 8 to 10 x 10^6^ bone marrow pooled cells from each genotype donors were transferred intravenously into lethally irradiated *Ldlr*^-/-^ recipient mice. Four weeks after transplantation, bone marrow reconstitution was checked in blood using the CD45 haplotypes as markers to detect donor and recipient cells.

### Single-Cell Sorting and Sequencing of Immunoglobulin Transcripts

Single cell sorting, and primer matrix-based Ig gene single cell PCR and sequencing were performed as previously described ^71,72^. Briefly, GC B cells (B220^+^ GL7^+^ tdTom^+^) and MBCs (B220^+^ GL7^+^ tdTom^+^) from lymph nodes and plasma cells PCs (CD138^+^tdTom^+^) from bone marrow were isolated from CFA and MDA-LDL+CFA immunized *Aicda*-Cre^+/ki^ *R26tdTom*^+/ki^ mice and single-cell sorted into 384-well plates filled with lysis buffer using an Aria III flow cytometric cell sorter and index sort option (BD Biosciences). RNA from single cells was retrotranscribed using random hexamer primers and cDNA was used as template for three independent nested PCRs with barcoded primers to amplify *Igh, Igk, Igl* transcripts. *Ighd*-specific primers were included in the Igh PCRs (1° PCR: mIghd-114-rv, 5’-CAGAGGGGAAGACATGTTCAACTAT-3’; 2° PCR: mIghd-079-rv, 5’-CAGTGGCTGACTTCCAATTACTAAAC-3’). Amplicons were pooled and sequenced on Illumina MiSeq 2<300 (Eurofins MWG) and data was processed and analyzed with sciReptor version v1.1-2-gf4cf8e2 ^73^.

### Enzyme-linked immunosorbent assay (ELISA) for immunoglobulin detection

Total, MDA-LDL- and MDA-BSA-specific IgG and IgM antibody titers in plasma or serum were determined using mouse IgM, mouse IgG, mouse IgG1, mouse IgG2b and mouse IgG2c ELISA Quantitation Kits or antibodies (Bethyl laboratories; E90-101, E90-131, A90-105P, A90-109P, A90-136P, respectively) used in accordance with the manufacturer’s instructions. In brief, plates were coated with goat anti-mouse IgM or IgG capture antibodies, MDA-LDL (3 μg/ml) or MDA-BSA (5 μg/ml) overnight at 4 °C. Then, plates were blocked and incubated with diluted plasmas (1/50 for IgM, 1/20 for IgG and 1/20000 for IgG1, IgG2b and IgG2c). Goat anti-mouse IgM, IgG, IgG1, IgG2b or IgG2c detection antibodies conjugated to HRP followed by TMB substrate solution (Bethyl laboratories, E102) were added to the plate and finally measure the absorbance at 450 nm. For competition immunoassays, pooled plasma (1/20000 dilution) from MDA-LDL+CFA immunized mice was preincubated with increasing concentrations (0, 25, 50, 100, 150, 200 and 300 μg/ml) of the indicated competitors (BSA, MDA-BSA, LDL and MDA-LDL) overnight at 4 °C. Plasma-competitor solutions were then centrifuged at 15.800 g for 45 minutes at 4 °C. Supernatants were added to MDA-LDL coated (3 μg/ml) ELISA plates after blocking. IgG1 bound antibodies from plasma were detected by adding a goat anti-mouse IgG1 detection antibody conjugated to HRP followed by the addition of TMB substrate solution. Results were expressed as ratios of IgG1 binding to MDA-LDL in the presence (B) or absence (B_0_) of the competitor.

### Quantification of atherosclerosis progression

Perfused hearts were fixed with 4% PFA 2 h at RT, incubated with 30% sucrose overnight, embedded in OCT compound (Sakura, 4583) and frozen using dry ice. Subsequently, aortic sinus was sectioned from proximal aorta to heart apex using a cryostat (10 μm sections). Atherosclerosis extent was evaluated by quantifying atherosclerotic plaque area in five to six serial 80 μm-spaced aortic sinus cross sections stained with Oil-Red-O. Mayer’s Hematoxylin (Bio-Optica, 05-06002/L) was used for nuclear counterstaining, and images were acquired with a DM2500 microscope (Leica). Image quantification was performed with Image J software ^74^. Area under the curve (AUC) analysis of quantified atherosclerosis value of Oil Red-O-stained serial 80 μm-spaced aortic sinus cryosections was performed using GraphPad Prism (version 7.03 for Windows, GraphPad Software, La Jolla California USA, www.graphpad.com). To normalize the AUC of *Prdm1*^+/+^ or *Prdm1*^fl/fl^ MDA-LDL+CFA and CFA groups to PBS, the AUC value of each mice was divided by the mean AUC of the PBS group with the same genotype (*Prdm1*^+/+^ or *Prdm1*^fl/fl^) and the result was multiplied by 100. Collagen content within atherosclerotic lesions was quantified from Picrosirius Red-stained cryosections and images were taken using ECLIPSE 90i microscope (Nikon). Plasma or serum lipid and lipoprotein (LDL cholesterol, HDL cholesterol, triglycerides and total and free cholesterol) profile from pooled or individual animals was measured using Dimension RxL Max Analyzer (Siemens).

### Immunofluorescence

For aortic sinus immunofluorescence, heart cryosections were incubated with 20 mM NH_4_Cl 10 minutes at RT. Then, sections were permeabilized with 0.5% Triton X-100 (Sigma-Aldrich, T9284-500ml) in PBS 30 minutes at RT and blocked with 2% BSA and 5% Donkey Serum (Sigma-Aldrich, D9663) in PBS during 1 h at RT. Subsequently, Avidin/Biotin blocking kit (Vector Labs, SP-2001) was used to block endogenous biotin in the sample. Samples were incubated with commercial primary antibodies SMA-1 (1A4, ThermoFisher Scientific) and MAC-2 (M3/38, Cedarlane (TebaBio)) overnight at 4 °C. After washing, Goat anti-rat-Alexa Fluor 568 and Streptavidin-Alexa Fluor 488 (ThermoFisher Scientific) were added to the sample and incubated for 1h at RT. Slides were mounted with Prolong Gold (Life Technologies, P36930) after DAPI (Sigma-Aldrich, D9542) staining and images were taken using a SPE confocal microscope (Leica). Image quantification was performed with Image J software.

### Statistical analysis

Data are presented as means ± standard deviation (SD). Statistical analysis was performed in GraphPad Prism (version 7.03 for Windows, GraphPad Software, La Jolla California USA, www.graphpad.com). Normality of the data was assessed by D’Agostino & Pearson normality test. For data coming from a Gaussian distribution, two-tailed unpaired Student t-test (p value ≤ 0..05 was considered significant) was used when comparing two experimental groups, one-way ANOVA was used when comparing three experimental groups (p value ≤ 0..05 was considered significant after correcting by two-stage linear step-up procedure of Benjamini, Krieger and Yekutieli test) and two-way ANOVA was used when comparing two experimental groups along time (p value ≤ 0.05 was considered significant after correcting by two-stage linear step-up procedure of Benjamini, Krieger and Yekutieli test). When analyzing not normally distributed data, Mann Whitney test was used for analyzing two groups (p value ≤ 0.05 was consider significant) and Kruskal–Wallis test (p value ≤ 0.05 was consider significant after correcting for multiple comparison by two-stage linear step-up procedure of Benjamini, Krieger and Yekutieli test) was used for analyzing three or more groups. Fisher Exact and Chi-Square statistical (Chi-square Calculator - Unlimited Contingency Table Size (icalcu.com)) tests were used for categorical data (p value ≤ 0.05 was consider significant).

## ACKNOWLEDGEMENTS

We thank all members of the B Cell Biology Laboratory for useful discussions; A. del Monte for helpful advice and discussion; V. G. de Yébenes for critical reading of the manuscript; V. Labrador for help with microscopy and image analysis. I.M.-F. and C.L. were fellow of the research training program funded by Ministerio de Economía y Competitividad (SVP-2014-068216 and SVP-2014-068289, respectively); and A.R.R. was supported by Centro Nacional de Investigaciones Cardiovasculares (CNIC). The project leading to these results has received funding from la Caixa Banking Foundation under the project code HR17-00247 and from SAF2016-75511-R and PID2019-106773RB-I00 grants to A.R.R. (Plan Estatal de Investigación Científica y Técnica y de Innovación 2013–2016 Programa Estatal de I+D+i Orientada a los Retos de la Sociedad Retos Investigación: Proyectos I+D+i 016, Ministerio de Economía, Industria y Competitividad) and co-funding by Fondo Europeo de Desarrollo Regional (FEDER). CIBERCV (J.L.M-V.) and CIBERDEM (J.C.E-G.) are Instituto de Salud Carlos III projects. The CNIC is supported by the Instituto de Salud Carlos III (ISCIII), the Ministerio de Ciencia e Innovación (MCIN) and the Pro CNIC Foundation and is a Severo Ochoa institute (CEX2020-001041-S grant funded by MCIN/AEI /10.13039/501100011033).

---

**Figure S1. MDA-LDL immunization of pro-atherogenic mice. (A)** Weight gain of CFA and MDA-LDL+CFA immunized mice along the experiment. Quantification of **(B)** spleen GC B cells (B220^+^GL7^+^) and **(C)** bone marrow PCs (B220^-^CD138) from MDA-LDL+CFA and CFA mice by FACS. Population percentages showed in FACS graphs are frequency of alive cells. Data analyzed by unpaired t and two-way ANOVA statistical tests.

**Figure S2. *Aicda*-Cre^+/ki^ R26tdTom^+/ki^ reporter mice.** Genetic approach to monitor the GC response. In this model, the cDNA coding for the tdTom fluorescent protein is inserted in the Rosa26 endogenous locus preceded by a transcriptional stop sequence that is flanked by loxP sites. The Cre recombinase is inserted in the endogenous Aicda locus. In *Aicda*-Cre^+/ki^ *R26tdTom*^+/ki^ mice, activation of the Aicda locus promotes Cre expression and excision of the transcriptional stop at the R26tdTom allele, unleashing the expression of the tdTom protein.

**Figure S3. The MDA-LDL antibody repertoire. (A)** Representative plots of single cell sorted populations from CFA and MDA-LDL+CFA mice. **(B)** Circos plot representation of the 597 Igs sequenced in this study for the indicated features. Clones were stablished as B cells sharing the same V(D)J rearrangement and CDR3 length in both IgH and IgL. Shared clones are those containing B cells from both group of mice. Cell population origin, CSR, immunization group and number of Ig mutations (SHM) (nucleotide exchanges) is indicated for each B cell.

**Figure S4. Analysis of VJ gene usage and CDR3 length of MDA-LDL antibody repertoire.** Data shown summarize the *Ig*H, *Ig*ĸ and Ig⍰ V and J segment usage of Igs from single GC, MBC and PC derived from CFA and MDA-LDL+CFA immunized mice: **(A)** VH, **(B)** JH, **(C)** Vĸ, **(D)** Jĸ, **(E)** Vλ and **(F)** Jλ segment usage. CDR3 amino acid number in **(G)** IgH and **(H)** IgL. P-values were calculated by Fisher’s exact and non-parametric Mann-Whitney statistical tests. *P≤0.05, **P<0.01 and ****P<0.0001.

**Figure S5. Isotype distribution and mutational load in MDA-LDL antibody repertoire.** Number of IgH somatic mutations (nucleotide exchanges) within mutated **(A)** IgD, **(B)** IgG2b, **(C)** IgG2c, **(D)** IgG3 and **(E)** IgA Igs in each group. Isotype distribution within each sorted B cell population: **(F)** GC, **(G)** MBC and **(H)** PC. P-values were calculated by Fisher’s exact and non-parametric Mann-Whitney statistical tests. **P<0.01.

**Figure S6. Analysis of atherosclerotic plaque composition of *Prdm1*^fl/fl^ *Aicda*-Cre^+/ki^ *Ldlr*^-/-^ chimeras. (A)** Weight gain of *Prdm1*^+/+^ *Aicda*-Cre^+/ki^ *Ldlr*^-/-^ and *Prdm1*^fl/fl^ *Aicda*-Cre^+/ki^ *Ldlr*^-/-^ chimeras along the experiment. **(B)** Sirius Red staining of aortic sinus cryosections and **(C)** quantification of plaque collagen content. The scale bar in images represents 128 μm. **(D)** Representative MAC-2 (red) and SMA-1 (green) immunofluorescence in aortic sinus cryosections from *Prdm1*^+/+^ and *Prdm1*^fl/fl^ *Aicda*-Cre^+/ki^ *Ldlr*^-/-^ chimeras. Quantification of **(E)** plaque macrophage content (% MAC-2^+^) and **(F)** plaque smooth muscle cell content (% SMA-1^+^). The scale bar in images represents 172 μm. Data analyzed by unpaired t and two-way ANOVA statistical tests.

**Figure S7. Weight gain of immunized *Prdm1*^+/+^ and *Prdm1*^fl/fl^ *Aicda*-Cre^+/ki^ *Ldlr*^-/-^ chimeras.** Weight gain *Prdm1*^+/+^ and *Prdm1*^fl/fl^ *Aicda*-Cre^+/ki^ *Ldlr*^-/-^ chimeras immunized with MDA-LDL+CFA, CFA, and PBS along the experiment. Data analyzed by two-way ANOVA statistical test.

**Figure S8. Changes in lipid profile of *Prdm1*^+/+^ and *Prdm1*^fl/fl^ *Aicda*-Cre^+/ki^ *Ldlr*^-/-^ chimeras after immunization.** Plasma levels of **(A, B)** total and **(C, D)** free cholesterol, **(E, F)** LDL cholesterol, **(G, H)** triglycerides and **(I, J)** HDLcholesterol in *Prdm1*^+/+^ *Aicda*-Cre^+/ki^ *Ldlr*^-/-^ **(A, C, E, G and I)** and *Prdm1*^fl/fl^ *Aicda*-Cre^+/ki^ *Ldlr*^-/-^ **(B, D, F, H and J)** chimeras after immunization with PBS, CFA, or MDA-LDL+CFA. Data analyzed by one-way ANOVA statistical test. *P≤0.05 and **P<0.01.

