## Supplementary figures and images for "MDA-LDL vaccination induces athero-protective germinal center-derived antibody responses"

### Supplementary Figures

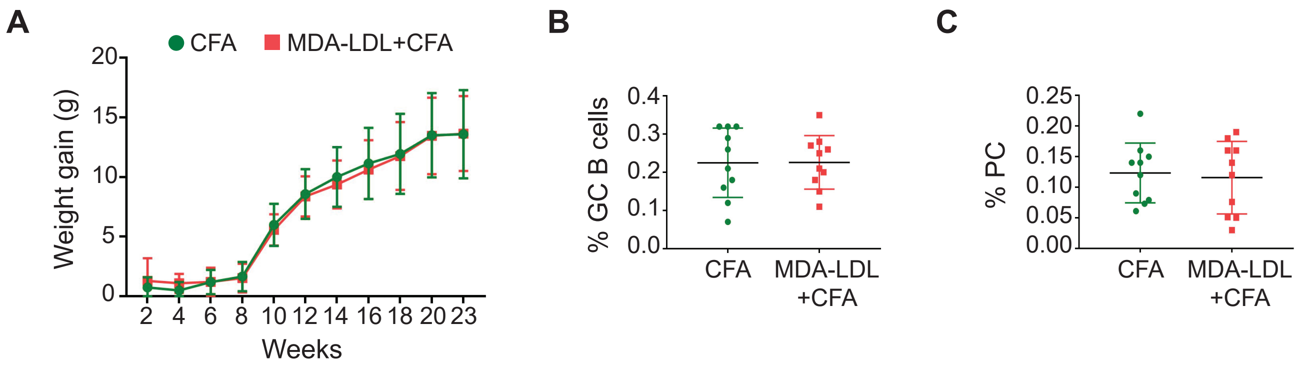

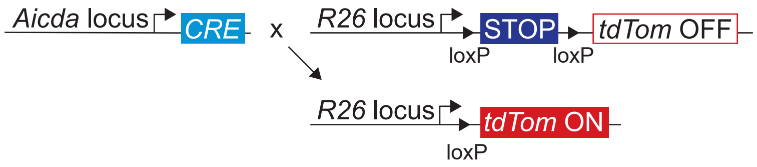

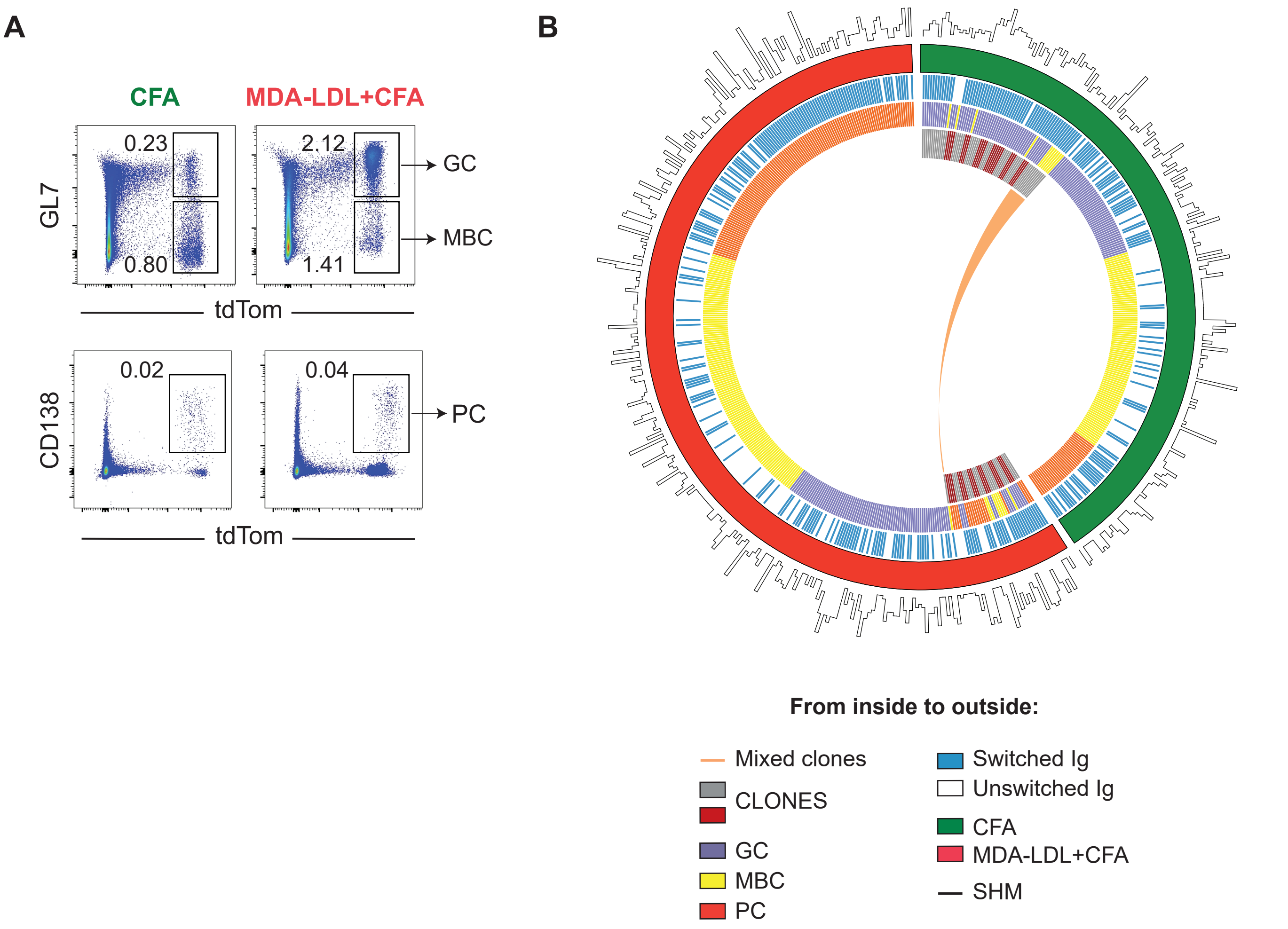

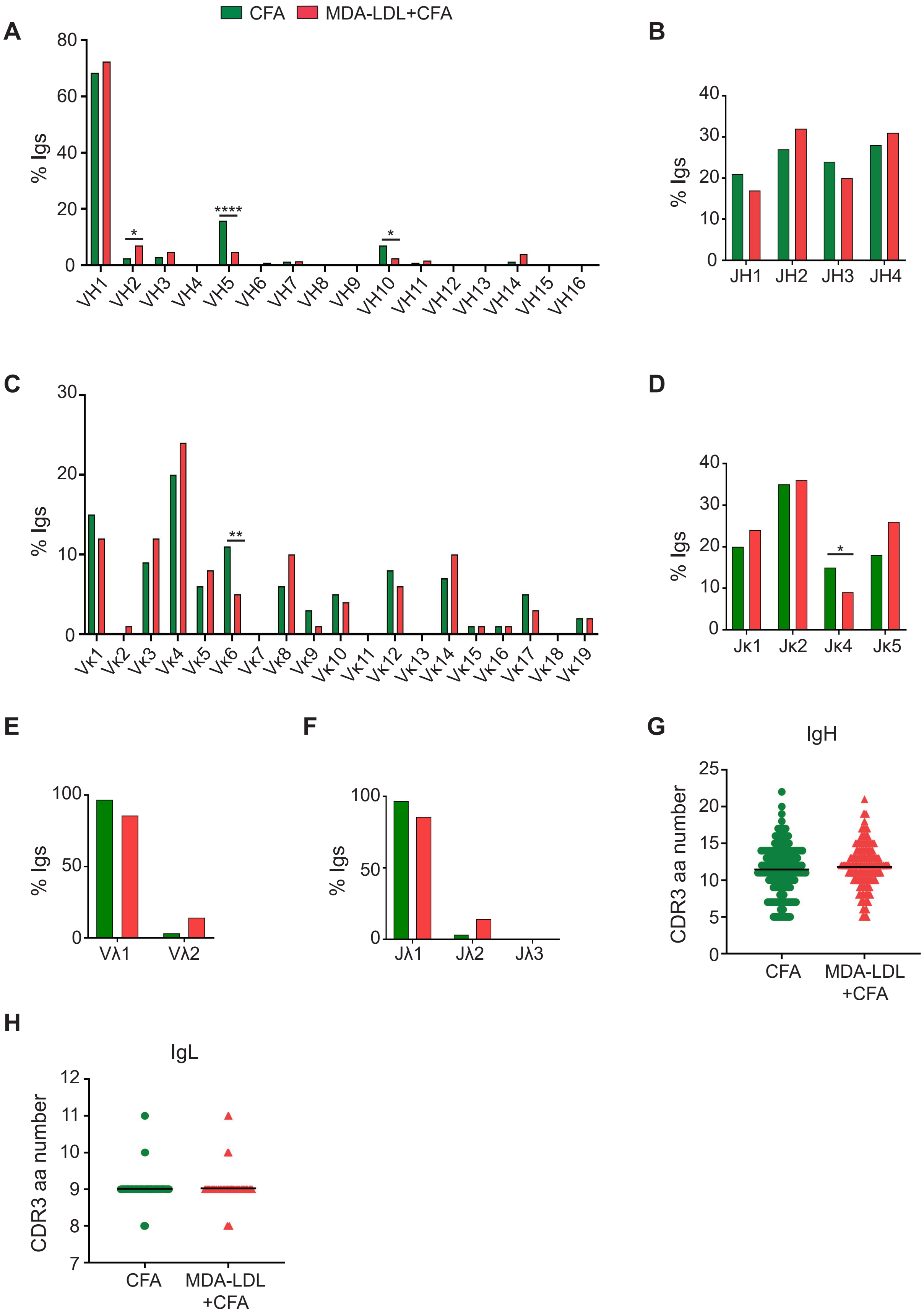

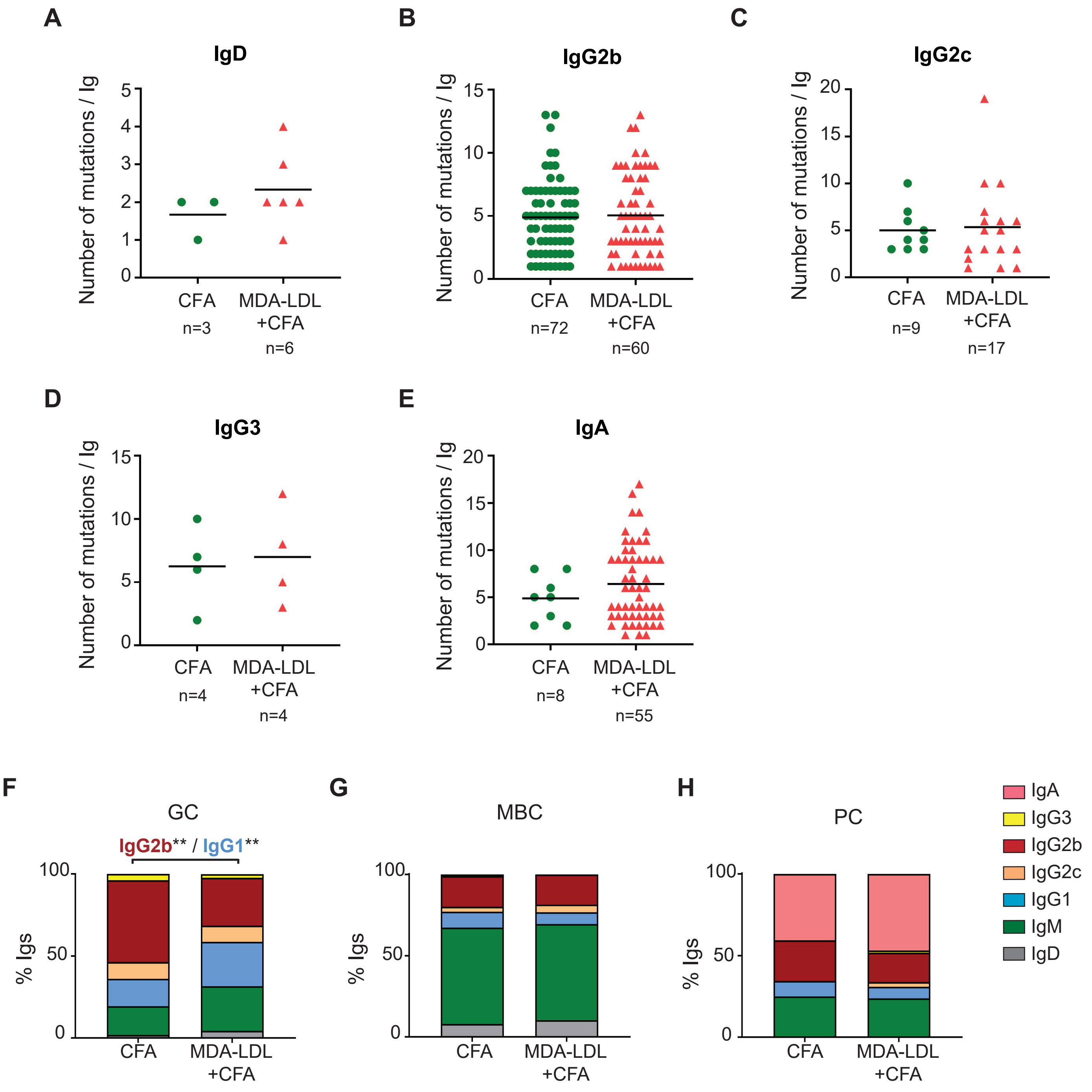

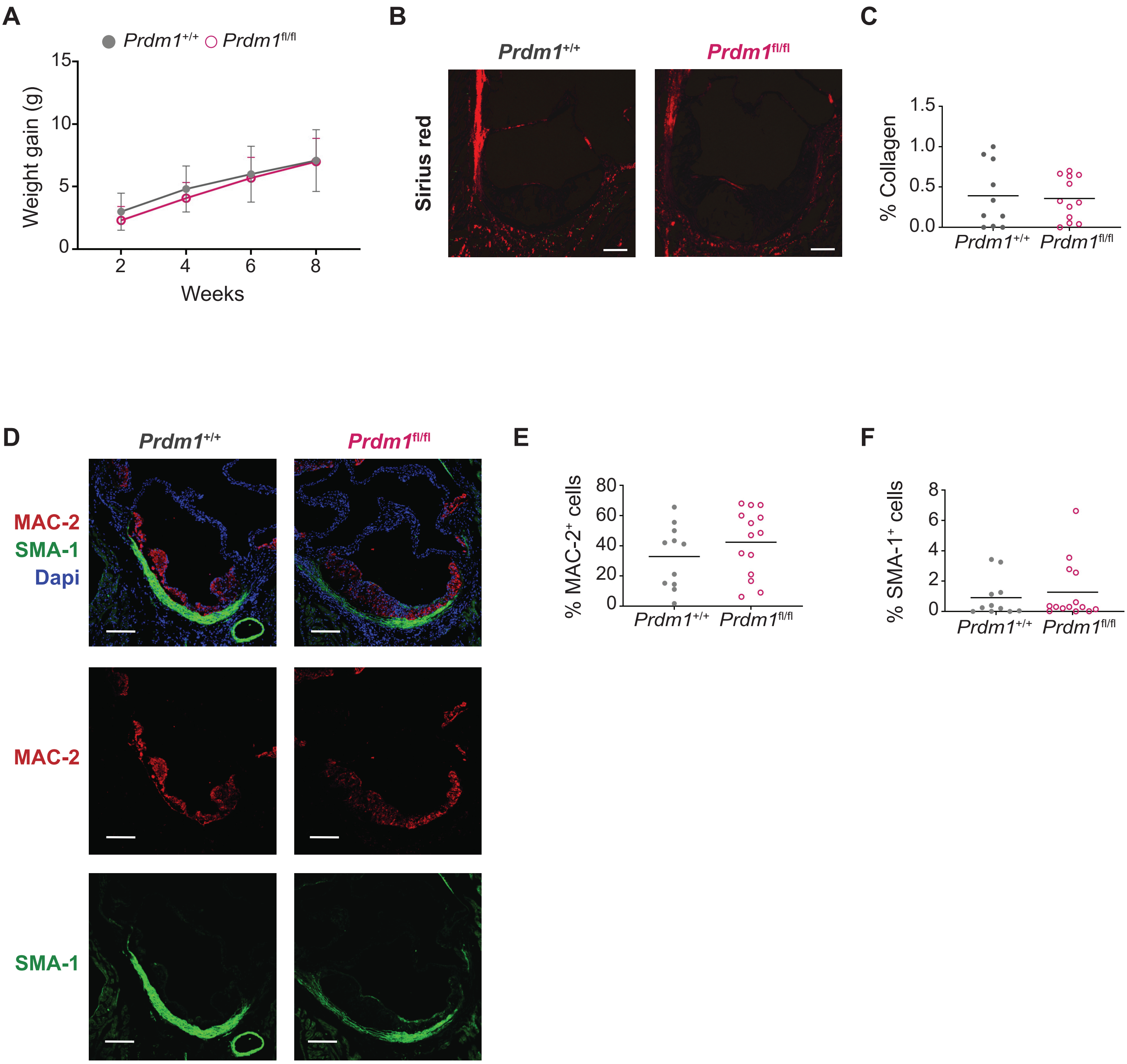

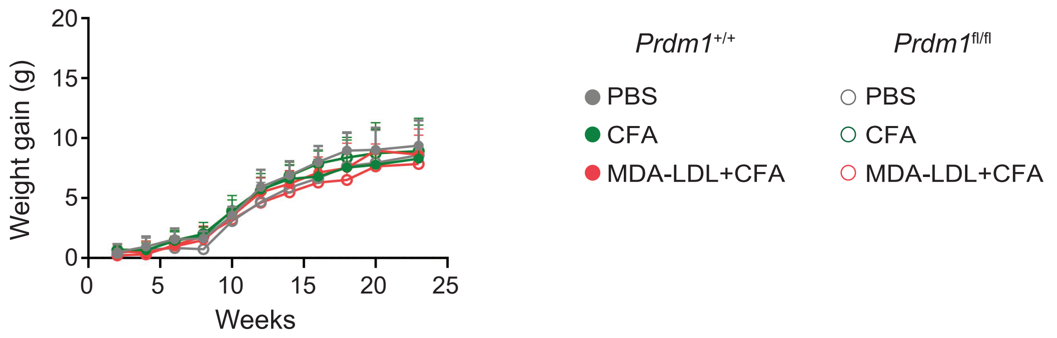

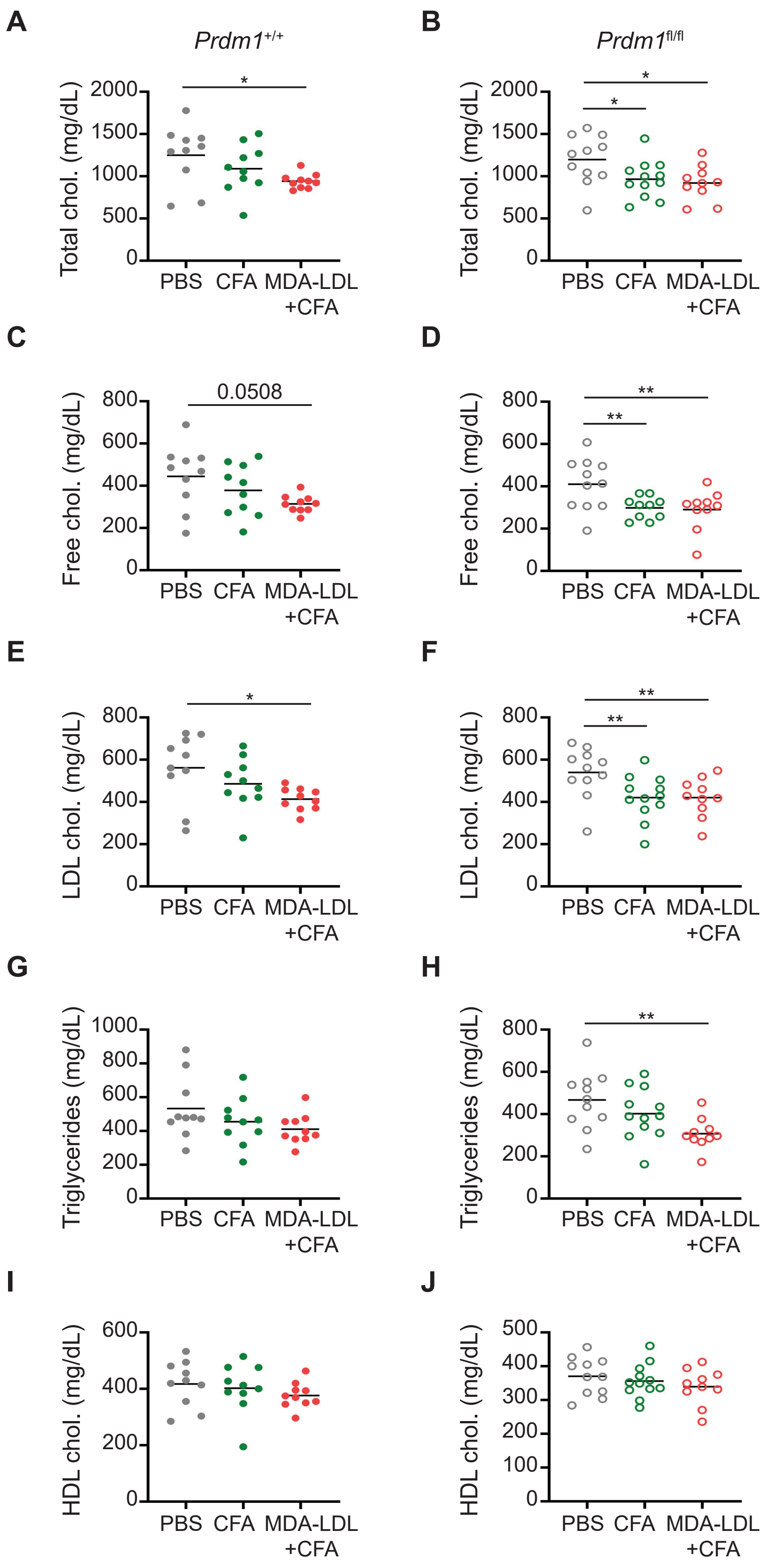
